# Symmetric Self-play Online Preference Optimization for Protein Inverse Folding

**DOI:** 10.64898/2026.03.26.714453

**Authors:** Wenwu Zeng, Xiaoyu Li, Haitao Zou, Yutao Dou, Xiongjun Zhao, Shaoliang Peng

## Abstract

Multi-objective reinforcement learning based on predicted structure feedback has been introduced into protein inverse folding. However, existing methods typically rely on a single model to optimize multiple structural objectives via a scalarized reward, which can bias the optimization toward dominant objectives and limit the exploration of diverse solutions. Here, we propose a online **Symmetric Self-play Preference Optimization (SSP)** framework that decouples the optimization of multiple structural objectives by training separate preference models with distinct reward signals, while enabling interaction through a shared sampling pool. This design allows the models to explore diverse optimization trajectories without enforcing a single dominant direction. Extensive experiments on both natural and *de novo* binder backbone inverse folding tasks demonstrate that SSP consistently improves sequence design self-consistency compared to single-model and existing baselines. Further analysis shows that different structural objectives are only partially aligned and induce distinct optimization directions, as evidenced by metric correlation and white-box analyses. This supports the effectiveness of decoupling objectives to enable higher design quality in protein design.

## 1 Introduction

Protein design has transformative potential across multiple areas of biotechnology, such as gene editing, large-molecule drug discovery, and immunotherapy [1, 2]. Existing AI-driven protein design methods mainly consist of three steps: **1**. *de novo* backbone generation (e.g. RFdiffusion [3]); **2**. inverse folding (IF) for sequence design (e.g. ProteinMPNN [4]); **3**. structure prediction (e.g. AlphaFold3 [5])/molecular dynamics (e.g. GROMACS [6]) screening. Among these protocol, protein IF constitutes a critical bottleneck. The amino acid sequence space grows exponentially with sequence length (*O*(*n*^20^)), while only a tiny subset of sequences can reliably fold into a specified structure. Consequently, IF models must effectively navigate an extremely high-dimensional search space to identify viable sequence solutions compatible with the target backbone.

Post-AlphaFold2 era [7], protein inverse folding (IF) has seen rapid progress, as accurate structure prediction enables the use of structural consistency as a evaluation signal. Early approaches such as ESM-IF [8], ProteinMPNN [4], and LigandMPNN [9] leverage geometry-aware neural networks to encode backbone structures and generate sequences via autoregressive/random order decoding. More recent works extend this paradigm through multimodal pretraining (e.g., ESM3 [10]), diffusion-based generation [11–13], and retrieval-augmented design [14, 15], further improving representation capacity and exploration ability.

Despite these advances, IF remains inherently underdetermined: multiple sequences can fold into similar structures, implying that a single optimal solution may not exist. Consequently, effective design requires not only structural fidelity but also sufficient exploration of the sequence space. To address this, recent works introduce reinforcement learning (RL) or direct preference optimization (DPO) [16], where structure-based metrics (e.g., TM-score or pTM) are used as feedback signals to guide sequence generation. Existing approaches typically fall into two categories. The first optimizes a single objective, encouraging the model to favor sequences with higher structural score/experimental stability [17, 18]. The second combines multiple objectives into a scalar reward, often via weighted aggregation [19], or constructs separate preference datasets for different metrics [20].

Considering the inherent multi-attribute characteristics of proteins, the multi-objective optimization thinking for protein IF is generally correct. Existing multi-objective RL-based IF methods approaches rely on a single model to balance multiple objectives. However, since different structural metrics are only partially aligned (see Figure 5b), collapsing them into a single objective can induce a dominant optimization direction, limiting diversity and potentially overlooking promising candidates. To address this issue, we propose the **Symmetric Self-play Preference Optimization (SSP)**, an online RL framework that explicitly models multiple objectives through interacting policies. Instead of forcing a single policy to balance heterogeneous signals, SSP maintains multiple policies that specialize in different objectives and learn collaboratively through a shared sampling mechanism. This design enables both diverse exploration and mutual enhancement, leading to improved design quality.

We implement the SSP framework on three representative sequence design models, including ESM3, ESM-IF1, and ProteinMPNN, demonstrating its architectural generality. Experimental results on widely used CATH benchmarks show that SSP consistently outperforms state-of-the-art (SOTA) methods in terms of self-consistency and structure prediction confidence. Meanwhile, evaluations on CAMEO targets with low structural similarity, as well as *de novo* binder design tasks across diverse biomolecular targets, demonstrate the robustness and transferability of SSP in challenging and out-of-distribution settings. Molecular dynamics simulations on DNA- and peptide-binding cases show that our designs not only possess high static structural fidelity and confidence, but also sustain stable interactions with their target biomolecules. Furthermore, diversity, novelty and correlation analysis showed that SSP concentrates sampling into high-quality structural regions while decoupling partially aligned objectives, enabling the discovery of high-fidelity, and novel protein sequences.

## 2 Preliminaries

### 2.1 Protein inverse folding

Protein IF aims to generate an amino acid sequence that folds into a given backbone structure. Formally, given a backbone structure **X** ∈ ℝ^*L*×4×3^ consisting of *L* residues (each represented by backbone atom coordinates), the goal is to generate a sequence **S** = (*s*_1_, *s*_2_, …, *s*_*L*_), where *s*_*i*_ denotes the amino acid type at position *i*.

#### Supervised inverse folding

Traditional inverse folding models are trained in a supervised manner by maximizing the conditional likelihood of the native sequence given the backbone structure:

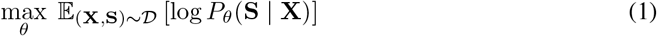

In practice, this is typically implemented using an autoregressive factorization:

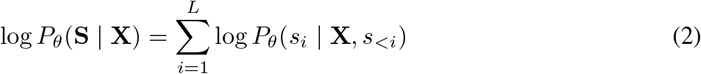

which encourages the model to recover native sequences conditioned on the backbone.

### 2.2 Reinforcement learning with structure feedback

Recent approaches incorporate RL to further improve sequence design by leveraging predicted structure-based feedback. Specifically, given a backbone **X**, a sequence **S** is sampled from the policy *π*_*θ*_(**S** | **X**), and a structure prediction model *f* is used to predict its folded structure 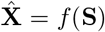. A reward function 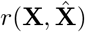 is then computed to measure structural quality (e.g., TM-score or pTM). The policy is optimized to favor sequences that yield higher rewards:

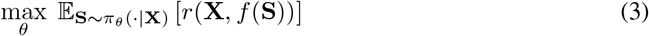

From the perspective of target rewards, existing RL-based methods for protein IF can be broadly categorized into two paradigms:

#### (1) Single-objective optimization

These methods optimize a single metric as the reward signal and construct preference pairs accordingly. Representative examples include DPO-based approaches such as ProteinDPO [17] and InstructPLM-DPO [18], which leverage experimental fitness and TM-score to guide sequence generation, respectively.

#### (2) Multi-objective optimization

Another line of work extends to multiple structural metrics. A common approach aggregates different objectives into a single scalar reward via weighted summation, as in ProteinZERO (TM-score + predicted free energy) [19], to balance various aspects of structural quality. Alternatively, methods such as MoMPNN [20] construct multiple sampling pools based on different property predictors, including designability metrics TM-score and developability metrics such as solubility, thermostability, and evolutionary plausibility. Preference pairs are then constructed within each objective to guide multi-objective optimization.

## 3 Methods

### 3.1 Symmetric Self-Play Online Preference Optimization

We propose a online SSP framework for protein IF (see Algorithm 1 and Figure 1), where two policy models are jointly optimized under complementary objectives. Specifically, the framework consists of two policy networks, *π*_*A*_ and *π*_*B*_, and a slowly updated reference model *π*_ref_. The two policies are designed to capture different aspects of protein quality: *π*_*A*_ is optimized toward structural self-consistency (*R*_*sc*_), while *π*_*B*_ focuses on predictive structural confidence (*R*_*pred*_). The reference model *π*_ref_ serves as a stable anchor to regularize preference optimization and is updated using an exponential moving average (EMA) of the two policies.

**Figure 1.**
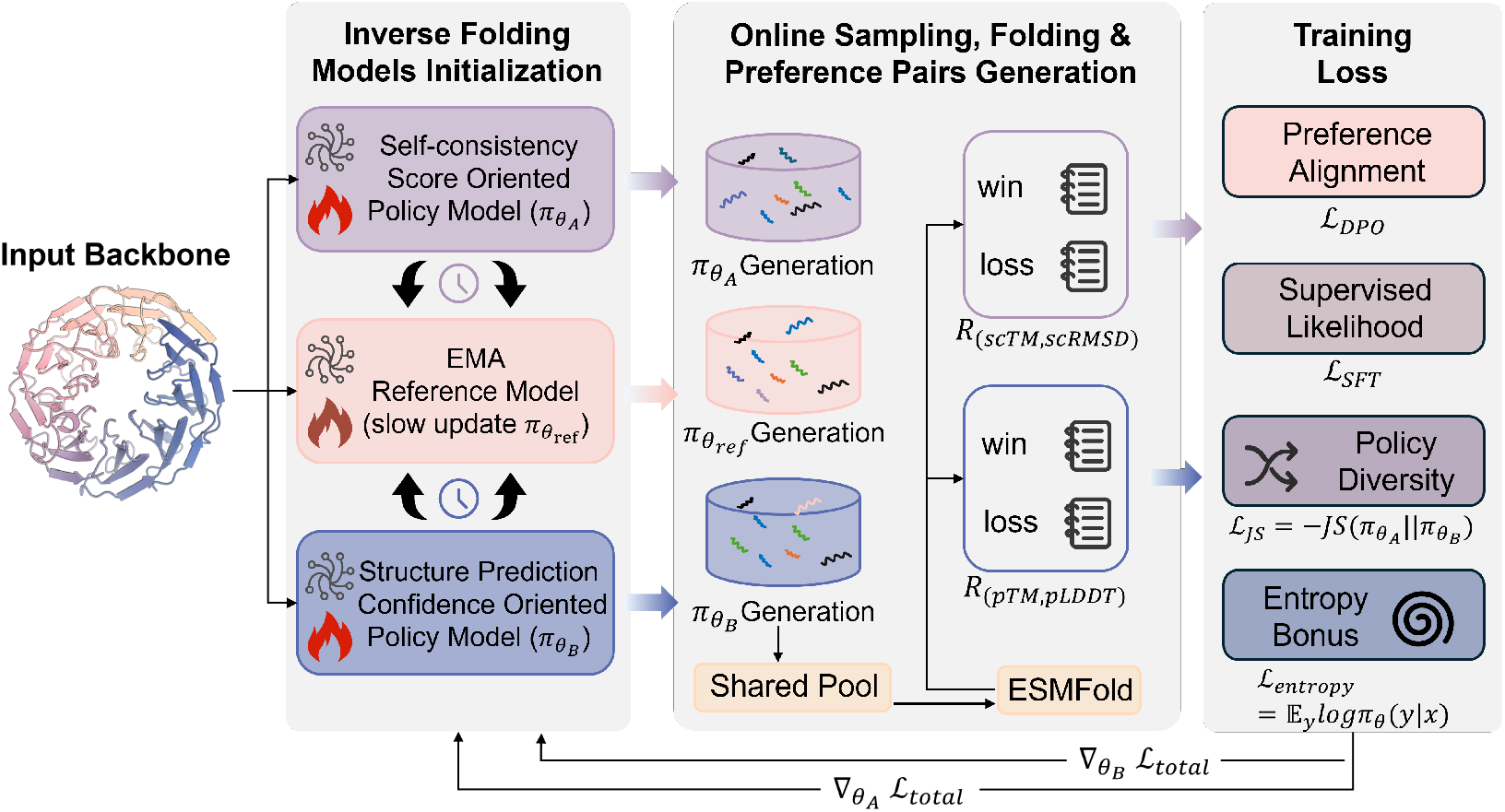
SSP framework.

Given a backbone structure *X*, each policy independently samples a set of candidate sequences. In our implementation, each model samples 5 sequences using temperature-controlled decoding (*T* = 1), and all candidates are merged into a shared pool: *Y* = *Y*_*A*_ ∪ *Y*_*B*_ ∪ *Y*_ref_. Each sequence *y* ∈ *Y* is refolded using ESMFold [21] to compute confidence metrics including *pLDDT* and *pTM*. To assess structural fidelity, we align the predicted structure with the target backbone *X* using USalign [22], obtaining *Cα-RMSD* and *scTM* scores. To ensure training quality, we discard low-confidence candidates whose *pLDDT*<0.45 or *pTM*<0.35. The remaining sequences are scored using two reward functions corresponding to the two objectives. Preference pairs are then constructed within the shared pool, enabling cross-policy comparison and implicit competition. This symmetric interaction encourages the two policies to explore different regions of the solution space while jointly improving overall quality.

To further consolidate the complementary behaviors learned by the two self-play agents, we introduce a model merging step that unifies *θ*_*A*_ and *θ*_*B*_ into a single deployable model. For full-parameter models such as ProteinMPNN, we adopt a task vector merging strategy. Specifically, we construct the merged model as

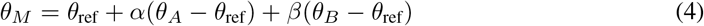

where *α* and *β* weight task-specific parameter shifts relative to the base model. This formulation allows us to linearly combine learned capabilities while preserving stability.

For parameter-efficient fine-tuning settings such as ESM-IF1 and ESM3, we instead merge Low-Rank Adaptation (LoRA) modules. Concretely, each LoRA module represents a low-rank update Δ*W* = *BA* applied to the base weights. Given two adapters, we compute their weighted combination:Δ*W*_*M*_ = *λ*_1_Δ*W*_*A*_ + *λ*_2_Δ*W*_*B*_, and inject the resulting update back into the base model.

The overall training loss function see the Appendix A.2. Implementation details and hyperparameters list see the Appendix A.7.

## 4 Results

We implemented the SSP framework on three IF models: ESM3 (∼1.4 billion parameters, multimodal pre-trained transformer architecture), ESM-IF1 (∼141 million parameters, GVP+autoregressive transformer architecture), and ProteinMPNN (∼1.66 million parameters, MPNN+random order decoding). ESM3 was trained on both CATH4.2 and CATH4.3 training sets respectively, while ESM-IF1 and ProteinMPNN were trained only once on the latter. Detailed information about all the datasets, metrics, and baseline methods in this study can be found at Appendixes A.3, A.4, and A.5.

### 4.1 Performance on native backbone benchmarks

We listed the performance comparisons with ESM-IF1, ProteinMPNN, ProteinDPO, InstructPLM [23], InstructPLM-DPO, ADFLIP, and MapDiff on the commonly used CATH4.2 and CATH4.3 test sets (see Tables 1 and 2). First, all of our SSP models show significant improvements compared to any fundamental model; Secondly, it achieved the best performance on both test sets compared to the SOTA method (including structure prediction metrics and self-consistency metrics); In particular, the SSP model still has advantages over classic DPO-based methods, namely ProteinDPO and InstructPLM-DPO. For example, when trained on ESM-IF1, the pTM and scTM of ESM-IF1_*pred*_ are 0.732 and 0.787, respectively, which are 0.68 and 0.89% higher than those of ProteinDPO, even though the latter used a larger training set and real experimental fitness as the reward.

**Table 1:**
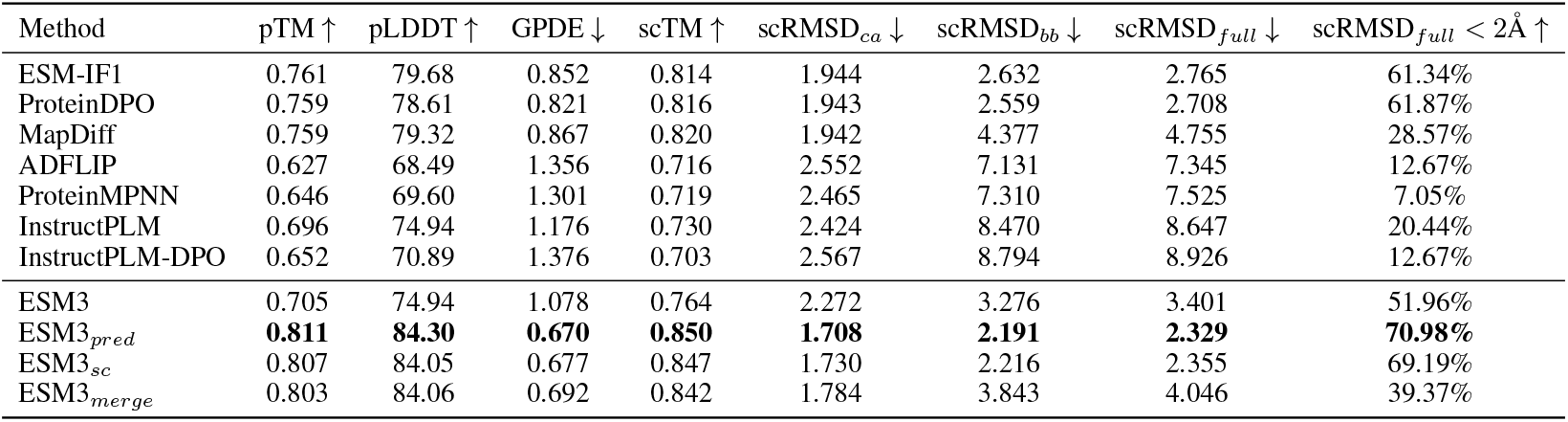
Performance on CATH4.2.

| Method | pTM ↑ | pLDDT ↑ | GPDE ↓ | scTM ↑ | scRMSD<sub>ca</sub> ↓ | scRMSD<sub>bb</sub> ↓ | scRMSD<sub>full</sub> ↓ | scRMSD<sub>full</sub> < 2Å ↑ |
| --- | --- | --- | --- | --- | --- | --- | --- | --- |
| ESM-IF1 | 0.761 | 79.68 | 0.852 | 0.814 | 1.944 | 2.632 | 2.765 | 61.34% |
| ProteinDPO | 0.759 | 78.61 | 0.821 | 0.816 | 1.943 | 2.559 | 2.708 | 61.87% |
| MapDiff | 0.759 | 79.32 | 0.867 | 0.820 | 1.942 | 4.377 | 4.755 | 28.57% |
| ADFLIP | 0.627 | 68.49 | 1.356 | 0.716 | 2.552 | 7.131 | 7.345 | 12.67% |
| ProteinMPNN | 0.646 | 69.60 | 1.301 | 0.719 | 2.465 | 7.310 | 7.525 | 7.05% |
| InstructPLM | 0.696 | 74.94 | 1.176 | 0.730 | 2.424 | 8.470 | 8.647 | 20.44% |
| InstructPLM-DPO | 0.652 | 70.89 | 1.376 | 0.703 | 2.567 | 8.794 | 8.926 | 12.67% |
| ESM3 | 0.705 | 74.94 | 1.078 | 0.764 | 2.272 | 3.276 | 3.401 | 51.96% |
| ESM3<sub>pred</sub> | **0.811** | **84.30** | **0.670** | **0.850** | **1.708** | **2.191** | **2.329** | **70.98%** |
| ESM3<sub>sc</sub> | 0.807 | 84.05 | 0.677 | 0.847 | 1.730 | 2.216 | 2.355 | 69.19% |
| ESM3<sub>merge</sub> | 0.803 | 84.06 | 0.692 | 0.842 | 1.784 | 3.843 | 4.046 | 39.37% |

**Table 2:** Performance on CATH4.3.

| Method | pTM ↑ | pLDDT ↑ | GPDE ↓ | scTM ↑ | scRMSD<sub>ca</sub> ↓ | scRMSD<sub>bb</sub> ↓ | scRMSD<sub>full</sub> ↓ | scRMSD<sub>full</sub> < 2Å ↑ |
| --- | --- | --- | --- | --- | --- | --- | --- | --- |
| ProteinDPO | 0.727 | 77.43 | 0.861 | 0.780 | 2.047 | 5.023 | 5.248 | 36.02% |
| MapDiff | 0.755 | 81.03 | 0.808 | 0.809 | 1.913 | 4.652 | 5.009 | 36.23% |
| ADFLIP | 0.591 | 67.71 | 1.389 | 0.669 | 2.696 | 7.837 | 8.054 | 10.52% |
| InstructPLM | 0.692 | 76.62 | 1.059 | 0.720 | 2.383 | 8.446 | 8.657 | 12.85% |
| InstructPLM-DPO | 0.650 | 73.03 | 1.253 | 0.693 | 2.550 | 8.850 | 9.029 | 11.53% |
| ProteinMPNN | 0.599 | 67.67 | 1.381 | 0.668 | 2.621 | 7.966 | 8.179 | 7.97% |
| ProteinMPNN<sub>pred</sub> | 0.694 | 75.90 | 1.006 | 0.737 | 2.199 | 6.733 | 6.937 | 15.14% |
| ProteinMPNN<sub>sc</sub> | 0.692 | 75.42 | 1.027 | 0.743 | 2.162 | 6.295 | 6.499 | 17.80% |
| ProteinMPNN<sub>merge</sub> | 0.698 | 75.97 | 1.005 | 0.744 | 2.147 | 6.269 | 6.468 | 20.51% |
| ESM-IF1 | 0.705 | 76.84 | 0.975 | 0.764 | 2.137 | 5.554 | 5.752 | 27.84% |
| ESM-IF1<sub>pred</sub> | 0.732 | 79.05 | 0.866 | 0.787 | 2.017 | 4.996 | 5.200 | **36.55%** |
| ESM-IF1<sub>sc</sub> | 0.730 | 78.85 | 0.889 | 0.785 | 2.019 | 5.116 | 5.298 | 28.79% |
| ESM-IF1<sub>merge</sub> | 0.727 | 78.64 | 0.885 | 0.782 | 2.054 | 5.149 | 5.351 | 28.58% |
| ESM3 | 0.697 | 75.93 | 1.009 | 0.758 | 2.210 | 5.576 | 5.781 | 26.24% |
| ESM3<sub>pred</sub> | 0.763 | 81.84 | 0.767 | 0.806 | 1.925 | 4.635 | 4.836 | 32.35% |
| ESM3<sub>sc</sub> | 0.753 | 81.01 | 0.797 | 0.801 | 1.953 | 4.697 | 4.901 | 31.56% |
| ESM3<sub>merge</sub> | **0.782** | **83.88** | **0.691** | **0.817** | **1.863** | **4.378** | **4.559** | 34.90% |

We further evaluated the generalization ability of the SSP model on CAMEO43, where the maximum TM-score of each samples against the CATH training set is less than 0.5 (see Table 3. All SSP models here are built on CATH4.3). First, compared to the three base models, the pTM values of the SSP merge model was improved by 13.90, 1.03, and 13.55% respectively; then, our best SSP model, ESM3_*merge*_, has pTM and pLDDT values of 0.762 and 82.68, respectively, which are 6.72 and 6.62% higher than those of the second-best method MapDiff. In terms of self-consistency metrics, ESM3_*sc*_ has a scTM of 0.774, which is 1.04% higher than that of Mapdiff. The results above demonstrate that: **1**. The SSP model has **model generalization ability**, which can be applied to different IF model architectures and improve backbone designability; **2. Data generalization ability**. Even on backbone with low structure similarity, it still outperforms SOTA methods in terms of structural prediction confidence and self-consistency.

**Table 3:** Performance on CAMEO43.

| Method | pTM ↑ | pLDDT ↑ | GPDE ↓ | scTM ↑ | scRMSD<sub>ca</sub> ↓ | scRMSD<sub>bb</sub> ↓ | scRMSD<sub>full</sub> ↓ | scRMSD<sub>full</sub> < 2Å ↑ |
| --- | --- | --- | --- | --- | --- | --- | --- | --- |
| InstructPLM | 0.593 | 69.20 | 1.526 | 0.605 | 3.185 | 8.903 | 8.988 | 20.93% |
| MapDiff | 0.714 | 77.54 | 0.989 | 0.766 | 2.355 | 3.524 | 3.715 | 44.18% |
| ADFLIP | 0.598 | 66.26 | 1.445 | 0.671 | 3.045 | 7.932 | 8.109 | 18.60% |
| ProteinDPO | 0.708 | 75.12 | 0.949 | 0.753 | 2.443 | 3.515 | 3.626 | 43.18% |
| InstructPLM-DPO | 0.638 | 70.10 | 1.398 | 0.667 | 3.114 | 9.594 | 9.601 | 9.30% |
| ProteinMPNN | 0.590 | 64.64 | 1.507 | 0.645 | 3.037 | 8.444 | 8.640 | 18.18% |
| ProteinMPNN<sub>pred</sub> | 0.698 | 74.74 | 1.028 | 0.747 | 2.466 | 6.734 | 6.855 | 20.45% |
| ProteinMPNN<sub>sc</sub> | 0.684 | 73.73 | 1.055 | 0.722 | 2.507 | 7.452 | 7.607 | 20.45% |
| ProteinMPNN<sub>merge</sub> | 0.670 | 72.53 | 1.110 | 0.717 | 2.629 | 6.778 | 6.946 | 22.72% |
| ESM-IF1 | 0.675 | 73.87 | 1.104 | 0.718 | 2.552 | 4.338 | 4.423 | 34.88% |
| ESM-IF1<sub>pred</sub> | 0.670 | 73.82 | 1.091 | 0.731 | 2.671 | 6.786 | 6.986 | 20.45% |
| ESM-IF1<sub>sc</sub> | 0.697 | 76.27 | 0.973 | 0.741 | 2.550 | 6.395 | 6.528 | 22.72% |
| ESM-IF1<sub>merge</sub> | 0.682 | 75.14 | 1.017 | 0.714 | 2.601 | 7.329 | 7.503 | 27.27% |
| ESM3 | 0.669 | 73.00 | 1.184 | 0.721 | 2.596 | 3.834 | 3.932 | 47.61% |
| ESM3<sub>pred</sub> | 0.732 | 78.86 | 0.902 | 0.773 | 2.304 | 3.320 | 3.467 | 47.61% |
| ESM3<sub>sc</sub> | 0.728 | 78.62 | 0.865 | **0.774** | 2.341 | **3.070** | **3.186** | **52.38%** |
| ESM3<sub>merge</sub> | **0.762** | **82.68** | **0.727** | 0.727 | **2.065** | 5.612 | 5.830 | 34.09% |

We also observed two phenomena: **1**. Improvement on ESM-IF1 is limited, not as good as Protein-MPNN and ESM3. Since the post-training and pre-training of ESM-IF1 share the same training set, CATH4.3, the search space under the same backbone conditions is extremely limited, making it difficult to further explore better solutions; **2**. Only the ESM3_*merge*_ model outperforms the single-biased model on CAMEO43 and CATH4.3. First, ESM3 has a larger number of parameters than the other two models, which allows it to explore more differentiated paths to the optimal solution. That is, it observes the backbone from different perspectives to design fitting sequences, and fusing these paths is obviously beneficial. Second, the design difficulty of samples in CATH4.3 (more complex and updated topology) and CAMEO43 (lower structural similarity) is greater than that of CATH4.2, and the two single-biased models can easily find similar optimization paths for easy samples, leading to fusion failure.

### 4.2 Performance on *de novo* biomolecular binder

Transferability on *de novo* artificial backbone is the gold standard for evaluating the potential of IF models to be applied to the real world. We constructed two *de novo* binder test sets using BoltzGen [24] and PXDesign [25]: BoltzGen-419 (containing 189 DNA-binding backbones, 77 RNA-binding backbones, and 153 peptide-binding backbones, with hotspots unspecified) and PXDesign-PPI226 (containing 226 protein-binding backbones, with hotspots specified.) The detailed construction protocols see Appendix A.3.

From Tables 4 and 5, ESM3_*merge*_ consistently achieved best performance across both test sets. For example, the pTM and scTM values of ESM3_*merge*_ on BoltzGen-419 are 0.727 and 0.838 respectively, which are 3.41 and 2.07% higher those of the second-best method ProteinDPO. Considering the design of complex interaction interfaces, the ipTM values of ESM3_*merge*_ on BoltzGen-419 and PXDesign-PPI226 are 0.348 and 0.267 separately, which achieves the improvements of 4.19 and 3.48% against the second-best methods MapDiff and ProteinDPO. In terms of design success rate, ESM3_*merge*_ is the only method that exceeds 70% on PXDesign-PPI226. Furthermore, the SSP algorithm also demonstrates universality on the other two base models. For example, ProteinMPNN_*merge*_ and ESM-IF1_*merge*_ achieved scTM scores of 0.742 and 0.733 on BoltzGen-419, which are 26.40 and 2.23% higher than those of vanilla ProteinMPNN and ESM-IF1, respectively.

**Table 4:**
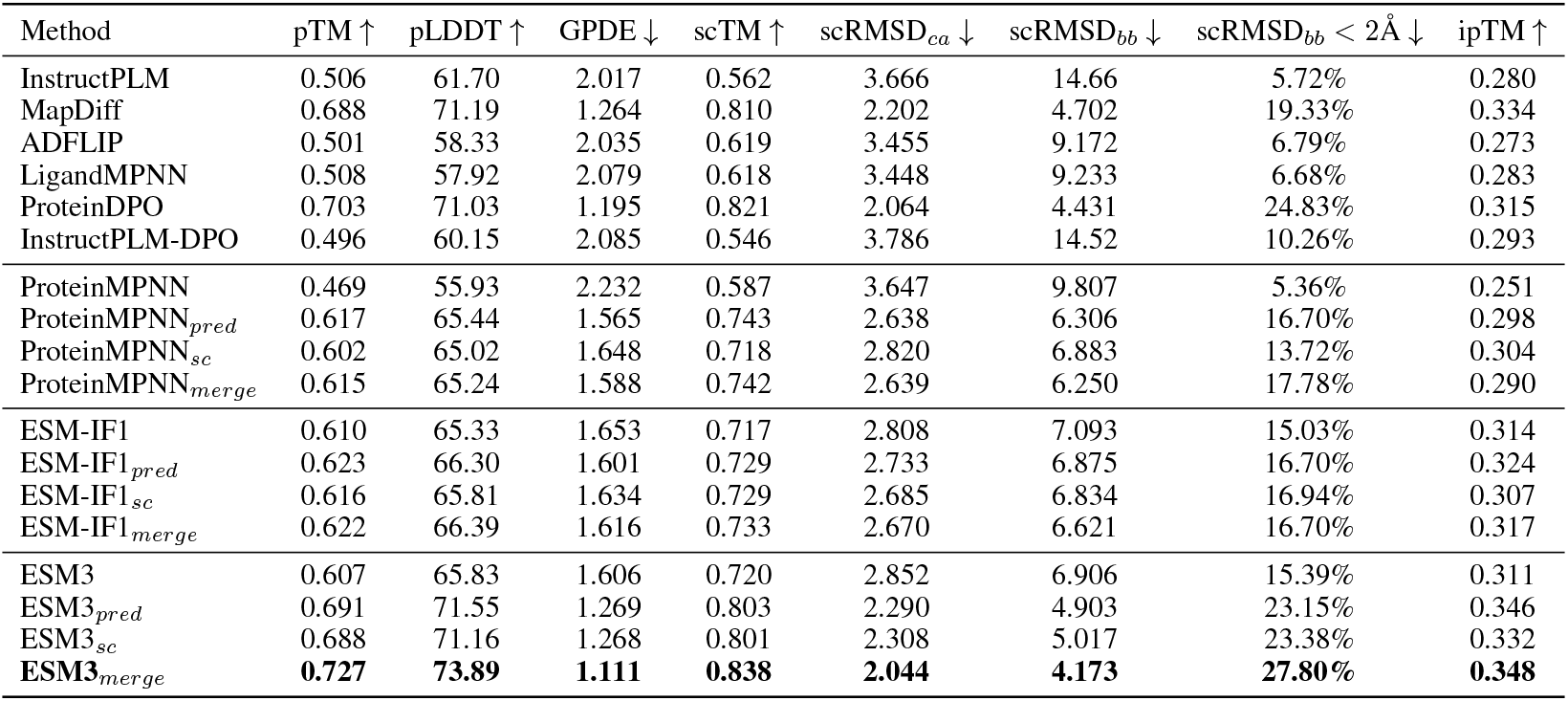
Performance on BoltzGen-419.

**Table 5:** Performance on PXDesign-PPI226.

| Method | pTM ↑ | pLDDT ↑ | GPDE ↓ | scTM ↑ | scRMSD<sub>ca</sub> ↓ | scRMSD<sub>bb</sub> ↓ | scRMSD<sub>bb</sub> < 2Å ↓ | ipTM ↑ |
| --- | --- | --- | --- | --- | --- | --- | --- | --- |
| InstructPLM | 0.528 | 66.97 | 3.085 | 0.486 | 3.339 | 18.12 | 0.00% | 0.184 |
| LigandMPNN | 0.592 | 71.47 | 1.391 | 0.582 | 2.982 | 8.263 | 22.12% | 0.254 |
| MapDiff | 0.569 | 69.91 | 1.734 | 0.560 | 3.113 | 9.351 | 7.52% | 0.219 |
| ADFLIP | 0.570 | 69.46 | 1.537 | 0.558 | 3.115 | 10.72 | 5.75% | 0.225 |
| ProteinDPO | 0.634 | 78.85 | 0.952 | 0.608 | 2.826 | 2.360 | 66.81% | 0.258 |
| InstructPLM-DPO | 0.524 | 65.18 | 3.202 | 0.489 | 3.358 | 17.784 | 0.00% | 0.197 |
| ProteinMPNN | 0.576 | 70.41 | 1.455 | 0.569 | 3.051 | 9.070 | 14.15% | 0.224 |
| ProteinMPNN<sub>pred</sub> | 0.614 | 76.49 | 1.120 | 0.590 | 2.874 | 4.853 | 46.01% | 0.261 |
| ProteinMPNN<sub>sc</sub> | 0.612 | 75.82 | 1.145 | 0.589 | 2.870 | 5.368 | 36.72% | 0.255 |
| ProteinMPNN<sub>merge</sub> | 0.610 | 76.15 | 1.132 | 0.592 | 2.855 | 4.926 | 43.80% | 0.243 |
| ESM-IF1 | 0.608 | 74.27 | 1.286 | 0.583 | 3.032 | 6.804 | 29.20% | 0.242 |
| ESM-IF1<sub>pred</sub> | 0.601 | 74.76 | 1.266 | 0.585 | 2.988 | 6.949 | 30.08% | 0.234 |
| ESM-IF1<sub>sc</sub> | 0.598 | 74.62 | 1.256 | 0.580 | 3.051 | 6.366 | 31.85% | 0.233 |
| ESM-IF1<sub>merge</sub> | 0.610 | 74.84 | 1.248 | 0.587 | 2.961 | 6.172 | 34.51% | 0.248 |
| ESM3 | 0.611 | 75.61 | 1.164 | 0.594 | 2.951 | 5.440 | 37.16% | 0.247 |
| ESM3<sub>pred</sub> | 0.629 | 78.58 | 1.041 | 0.602 | 2.851 | 3.271 | 57.52% | 0.263 |
| ESM3<sub>sc</sub> | 0.630 | 78.40 | 1.033 | 0.602 | 2.829 | 3.465 | 59.73% | 0.260 |
| ESM3<sub>merge</sub> | **0.640** | **80.75** | **0.923** | **0.610** | **2.703** | **2.248** | **70.35%** | **0.267** |

From the results above, the SSP framework demonstrates **transportability on the artificial *de novo* binder backbone** across different IF architectures and target biomolecules types. This characteristic produces a post-trained IF model, i.e., ESM3_*merge*_, that outperforms SOTA IF methods in preliminary computational evaluations.

### 4.3 Case study

Although structure prediction models can statically assess the designability of generated sequences, proteins require dynamic interactions with different biomolecules to perform their biological functions. Here, one *de novo* DNA-binding backbone (target PDB ID: 6ysl, chain C, BoltzGen generation) and one *de novo* peptide-binding backbone (target PDB ID: 8flj, chain M, BoltzGen generation) are employed for case studies. We first designed amino acid sequences using ESM3_*pred*_ and three SOTA sequence design methods: ESM3, LigandMPNN, and Mapdiff; the designed sequences were then input into the `AlphaFold3` online server ^3^ along with the target chain to predict the complex structure; finally, a 100 ns all-atom MD simulation was performed using `GROMACS` (see Appendix A.6 in detail).

From Figure 2, ESM3_*pred*_ consistently produces more reliable and dynamically stable complexes than competing methods. For the DNA-binding case (8flj), ESM3_*pred*_ achieves the best overall structural quality, with higher pTM/ipTM and more favorable binding energy. Importantly, MD simulations show that ESM3_*pred*_ maintains a stable complex throughout 100 ns, while all baselines exhibit substantial structural drift, indicating unstable protein–DNA interactions.

**Figure 2.**
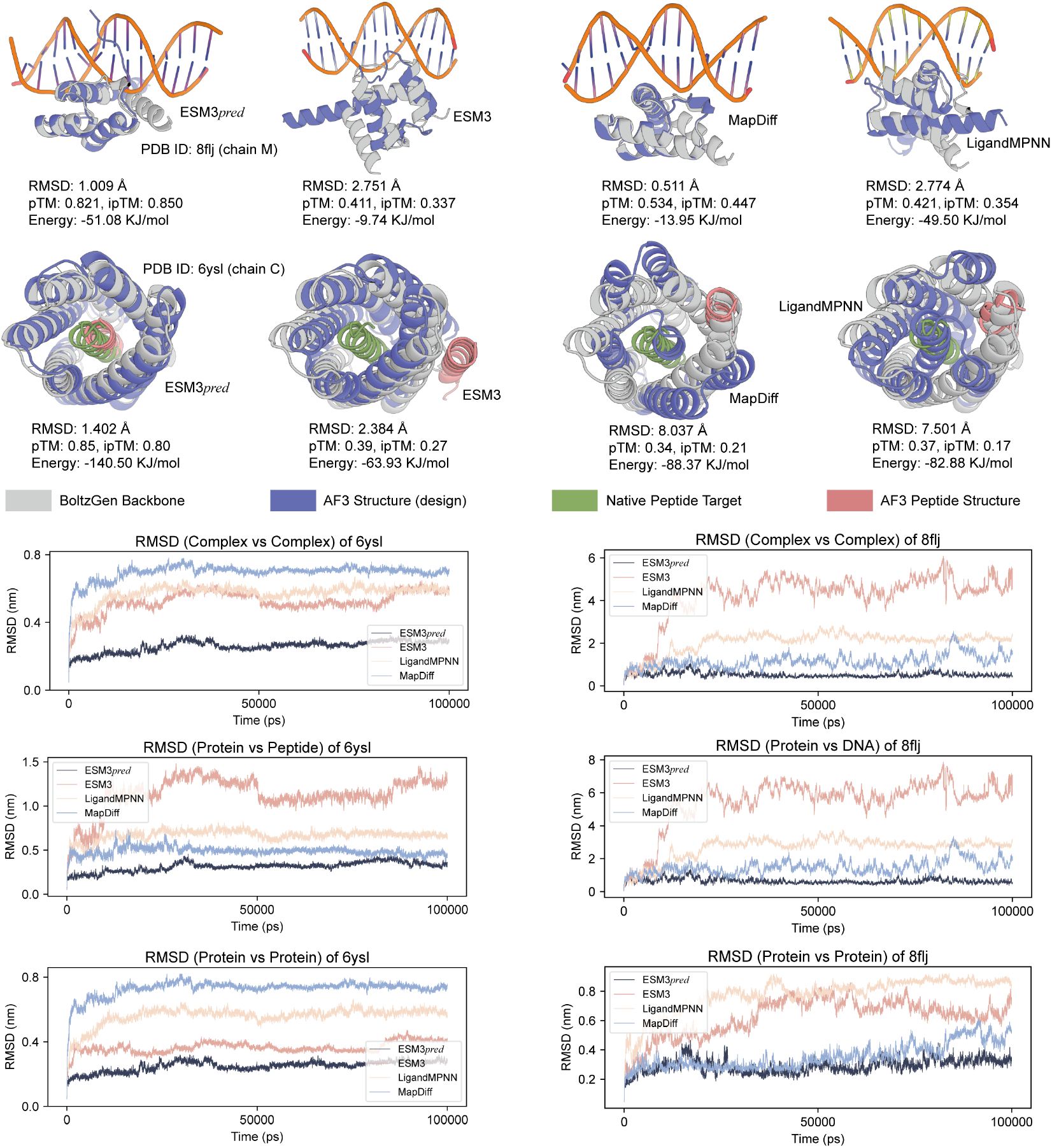
MD simulation on two cases, i.e., one DNA-binding protein (PDB ID: 6ysl, chain C) and one peptide-binding protein (PDB ID: 8flj, chain M). The free energy ↓ is calculated using the `gmx_MMPBSA` tool [26]. The RMSD curve depicts the amplitude of the dynamic motion of biomolecules system.

For the peptide-binding case (6ysl), a key difference lies in the recovered binding mode. Given that the binder backbone is *de novo* generated by BoltzGen, there is no true native complex to recover. Instead, we assess whether the designed sequence can realize a structurally consistent and physically plausible interaction pattern conditioned on the backbone. ESM3_*pred*_ successfully reconstructs a compact binding topology in which the backbone wraps around the target peptide, forming an enclosed interface as intended by the backbone geometry. In contrast, other methods fail to realize this enclosing configuration, leading to partially exposed or misaligned peptide placements. This discrepancy further results in reduced binding stability, as evidenced by higher RMSD and less favorable energies during MD simulations.

Overall, these results suggest that ESM3_*pred*_ not only improves global structure metrics but, more importantly, better captures the fine-grained geometric constraints required for stable biomolecular interactions.

### 4.4 Ablation study

To demonstrate the advantages of the dual-model self-play in our SSP architecture, we performed an ablation experiment on CATH4.2 (see Figure 3). Specifically, one policy model is removed from the SSP framework, retaining only a reference model and a policy model, which favors a single scTM (pTM or their weighted sum 0.5*scTM* + 0.5*pTM*). Consequently, we performed the same training protocol as SSP on ESM3, resulting in SP_*sc*_, SP_*pred*_, and SP_*mix*_. While the weighted fusion (like ProteinZERO [19]) of self-consistency and structural prediction confidence outperforms the single-biased model (like InstructPLM-DPO [18]), the improvement is limited. Contrarily, compared to the single-model DPO architecture, the SSP architecture has significant advantages over all metrics. Despite using the same base model, training data, and configuration as SP, SSP encourages exploration of diverse optimization directions. Coupled with the mutual enhancement induced by shared sampling pools, this design leads to consistent improvements over simple weight averaging and related baselines.

**Figure 3.**
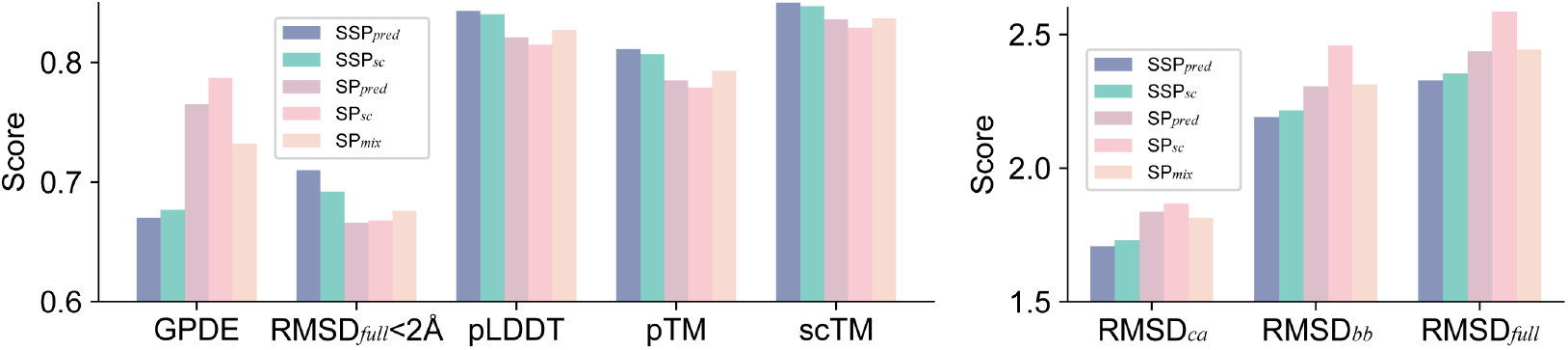
Ablation of SSP and single policy model (SP) architectures.

### 4.5 White-box Analysis of LoRA Update Geometry

Although the proposed SSP framework consistently improves both scTM and pTM scores on native and *de novo* binder backbone, it remains unclear whether these gains arise from redundant optimization or from genuinely diverse model behaviors. In particular, we aim to understand whether the scTM-optimized and pTM-optimized models learn similar parameter updates or explore different regions in parameter space. This distinction is crucial: if both models converge to similar update directions, the dual-model framework would be unnecessary; conversely, if they diverge, it would suggest that SSP enables complementary exploration.

To investigate this, we perform a white-box analysis of the learned LoRA updates for ESM3_*pred*_ and ESM3_*sc*_. For each LoRA layer, we extract the weight updates Δ*W*^*sc*^ = *B*^*sc*^*A*^*sc*^ and Δ*W*^*pred*^ = *B*^*pred*^*A*^*pred*^. We then compare the two updates using three complementary metrics. First, we measure the *subspace overlap* based on singular value decomposition (SVD), defined as Overlap = 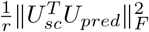, where *U*_*sc*_ and *U*_*pred*_ denote the left singular vectors of Δ*W*^*sc*^ and Δ*W*^*pred*^, respectively, and *r* is the LoRA rank. Second, we compute the *update direction similarity* via cosine similarity between flattened updates, 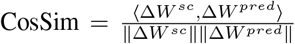. Finally, we compare the *update magnitudes* ∥Δ*W* ∥ to ensure that any observed differences are not due to scale discrepancies.

Our results reveal three key observations (see Figure 4). First, the subspace overlap remains consistently low across most layers (typically in the range of 0.05–0.15), indicating that the two models operate in largely distinct low-rank subspaces. Second, the cosine similarity between update directions is close to zero in most layers, suggesting that the parameter updates are nearly orthogonal and do not share a dominant direction. Third, the update magnitudes of the two models are comparable, ruling out trivial explanations based on scale differences. Interestingly, we observe a mild increase in both overlap and cosine similarity in deeper layers, implying partial alignment near the output space.

**Figure 4.**
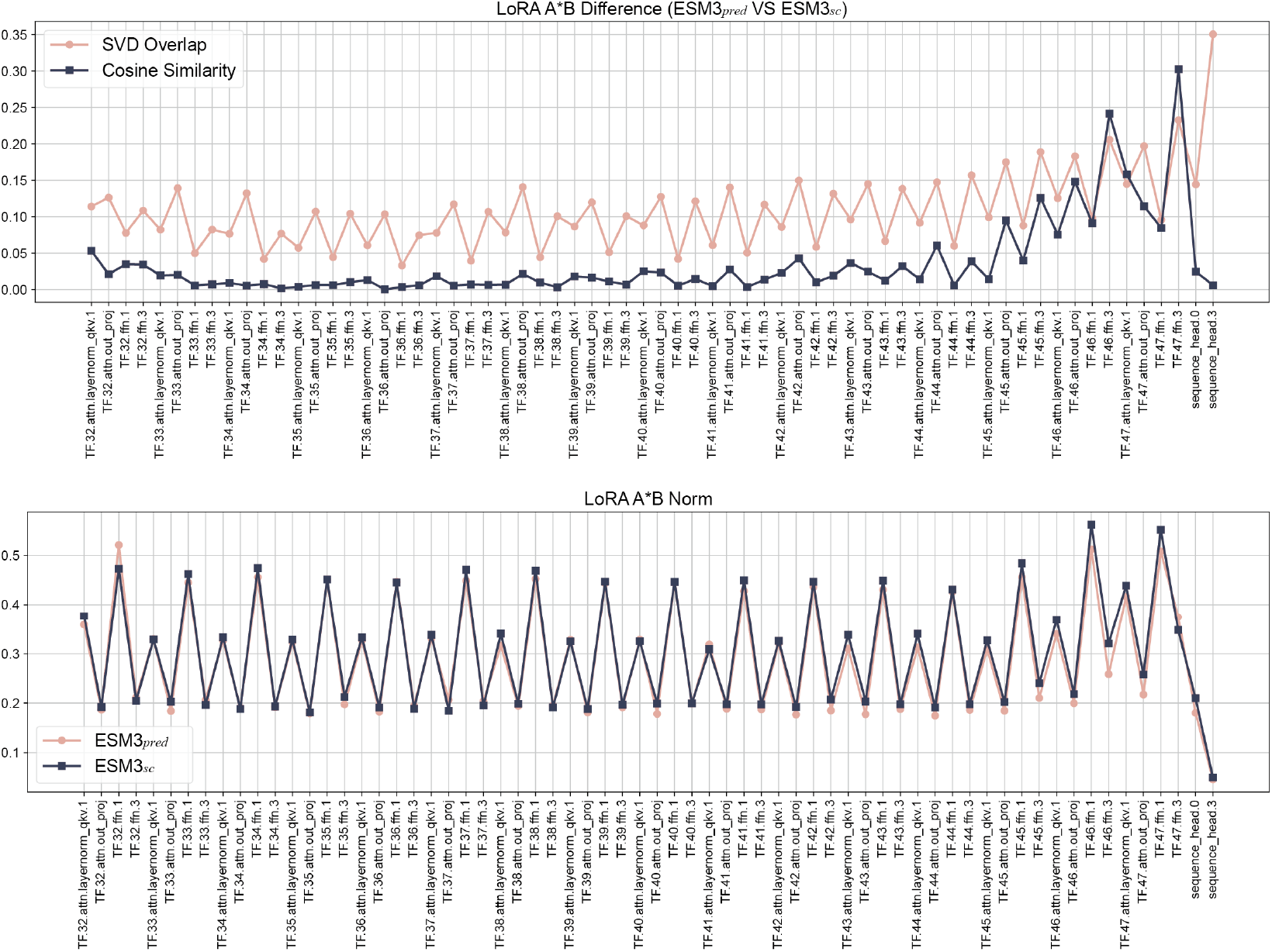
Model Intrinsic Analysis. The *sc*- and *pred*-biased models in SSP framework show low subspace overlap and weak directional alignment, despite having similar update magnitudes.

**Figure 5.**
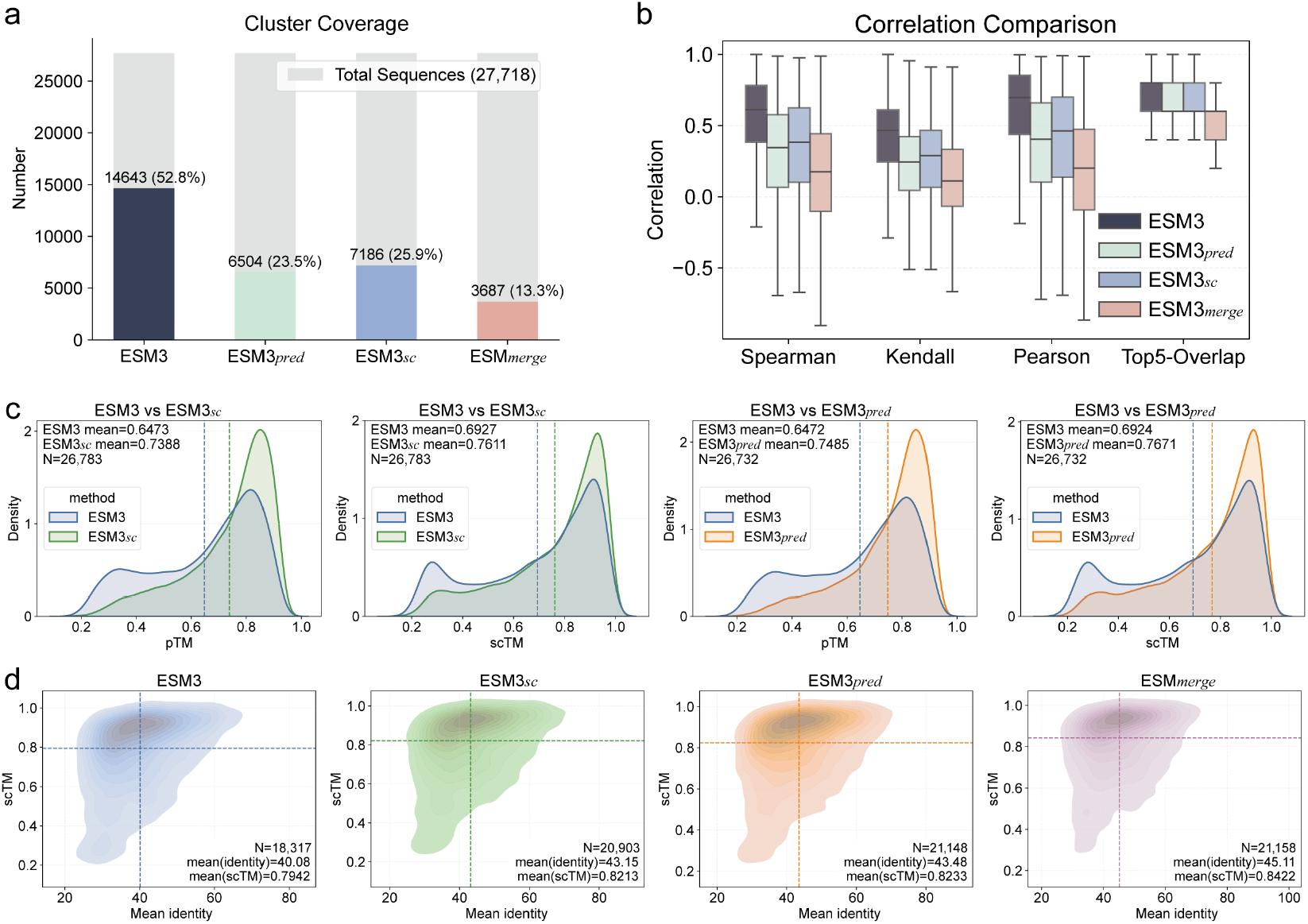
Diversity, novelty and relevance analysis.

Taken together, these findings suggest that optimizing for scTM and pTM leads to partially distinct parameter update geometries, with shared components but notable differences in the explored subspaces. Rather than being fully aligned or completely independent, the two models exhibit complementary update patterns in parameter space.

### 4.6 Diversity, novelty, and correlation analysis

To further investigate the effect of SSP on sequence design, we conduct a comprehensive analysis on the combined CATH4.2 and CATH4.3 test sets. For each backbone, we sample 10 sequences from different models, followed by structure prediction using Protenix. Structural quality is evaluated using pTM and scTM scores. In addition, sequence diversity is assessed by computing the average sequence identity against the matched sequences in UniRef50 and BFD Small databases using `DIAMOND blastp` [27].

#### SSP concentrates the solution space into high-quality structural regions

As shown in Figure 5a, the cluster coverage of SSP-based models is significantly reduced compared to the base model, indicating lower sequence-level diversity. However, this reduction does not imply a loss of effective solutions. Instead, combined with the distributions of pTM and scTM (Figure 5b) and the joint density plots over sequence identity and structural scores (Figure 5c), it is evident that SSP shifts the sampling distribution toward regions with higher structural quality. Specifically, both ESM3_*pred*_ and ESM3_*sc*_ exhibit a clear rightward shift in pTM and scTM, while the high-density regions in the identity–scTM space become more concentrated around high-scoring areas. This indicates that SSP filters out low-quality sequences and focuses exploration within a smaller but more reliable subset of the solution space.

#### SSP decouples partially aligned structural objectives and diversifies optimization directions

As illustrated in Figure 5b, the base model exhibits relatively high correlation between pTM and scTM, suggesting that these objectives are implicitly coupled during generation. In contrast, SSP significantly reduces this correlation across multiple metrics (Spearman, Kendall, and Pearson), while maintaining a relatively high top-k overlap. This indicates that SSP preserves coarse agreement among high-quality candidates, but introduces variability in their fine-grained ranking. In other words, SSP allows different policies to prioritize distinct structural aspects, resulting in a more diverse set of high-quality solutions across partially aligned objectives, rather than collapsing them into a single dominant optimization direction.

#### SSP generates sequences with both high self-consistency and high novelty

As shown in the bottom row of Figure 5d, SSP-based models achieve higher structural consistency while maintaining lower sequence similarity to known proteins. Specifically, both ESM3_*sc*_ and ESM3_*pred*_ exhibit improved scTM compared to the base model, and the merged model further enhances this trend, achieving the highest scTM (0.8422) among all variants. Importantly, the joint density distributions reveal that high-scoring regions (scTM > 0.8) are populated by sequences with relatively low identity, demonstrating that SSP is able to discover structurally consistent yet novel solutions. This suggests that SSP effectively breaks the conventional trade-off between structural fidelity and sequence novelty, enabling the generation of previously unseen sequences that remain compatible with the target backbone.

## 5 Conclusion

In this work, we study multi-objective reinforcement learning for protein inverse folding and show that partially aligned structural objectives can induce distinct optimization directions. To address the limitations of scalarized optimization, we propose a SSP framework that decouples objective optimization via separate preference models while enabling interaction through a shared sampling pool. This design promotes diverse optimization trajectories without enforcing a single dominant direction. Experiments demonstrate consistent improvements over single-model and existing baselines, particularly in challenging *de novo* settings.

Despite these promising results, several limitations remain. We outline several directions for future work in protein design. First, more effective co-optimization of physical objectives (e.g., stability and energetics) and structure-based metrics is needed to better align designed sequences with biophysical constraints. Second, incorporating realistic cellular environments, such as crowding effects and context-dependent interactions, could provide more faithful guidance for functional design. Third, developing more robust screening strategies to mitigate the limitations of current structure predictors, particularly for orphan proteins. Addressing these challenges would further improve the reliability and applicability of multi-objective protein design frameworks.

## 6 Data Availability

The data and source code of this study are freely accessible at https://github.com/wwzll123/SSP.

## 7 Author Contributions

Wenwu Zeng: Conceptualization, Methodology, Software, Visualization, Writing original draft. Xiaoyu Li: Visualization. Haitao Zou: Methodology. Yutao Dou: Software. Xiongjun Zhao: Conceptualization, Writing-review & editing. Shaoliang Peng: Funding acquisition, Resources, Supervision.

## 8 Acknowledgment

This work was supported by NSFC Grants 625B2068; NSFC-FDCT Grants 62361166662; National Key R&D Program of China 2023YFC3503400, 2022YFC3400400; The Innovative Research Group Project of Hunan Province 2024JJ1002; Hunan Science and Technology Innovation Plan 2025ZYJ003; Key R&D Program of Hunan Province 2023GK2004, 2023SK2059, 2023SK2060; Top 10 Technical Key Project in Hunan Province 2023GK1010; Key Technologies R&D Program of Guangdong Province (2023B1111030004 to FFH); Postgraduate Research Innovation Project of Hunan Province CX20250637. We would like to thank the Fund of the National Supercomputing Center in Changsha (http://nscc.hnu.edu.cn/), Peng Cheng Lab, Key Laboratory of High-Performance Distributed Ledger Technology and Digital Finance (Ministry of Education), and Hunan Research Center of the Basic Discipline for Cell Signaling.

## 9 Competing Interests

The authors declare no competing interests.

## A Supplementary Material

### A.1 SSP Algorithm

#### Algorithm 1 SSP Training for Protein Inverse Folding

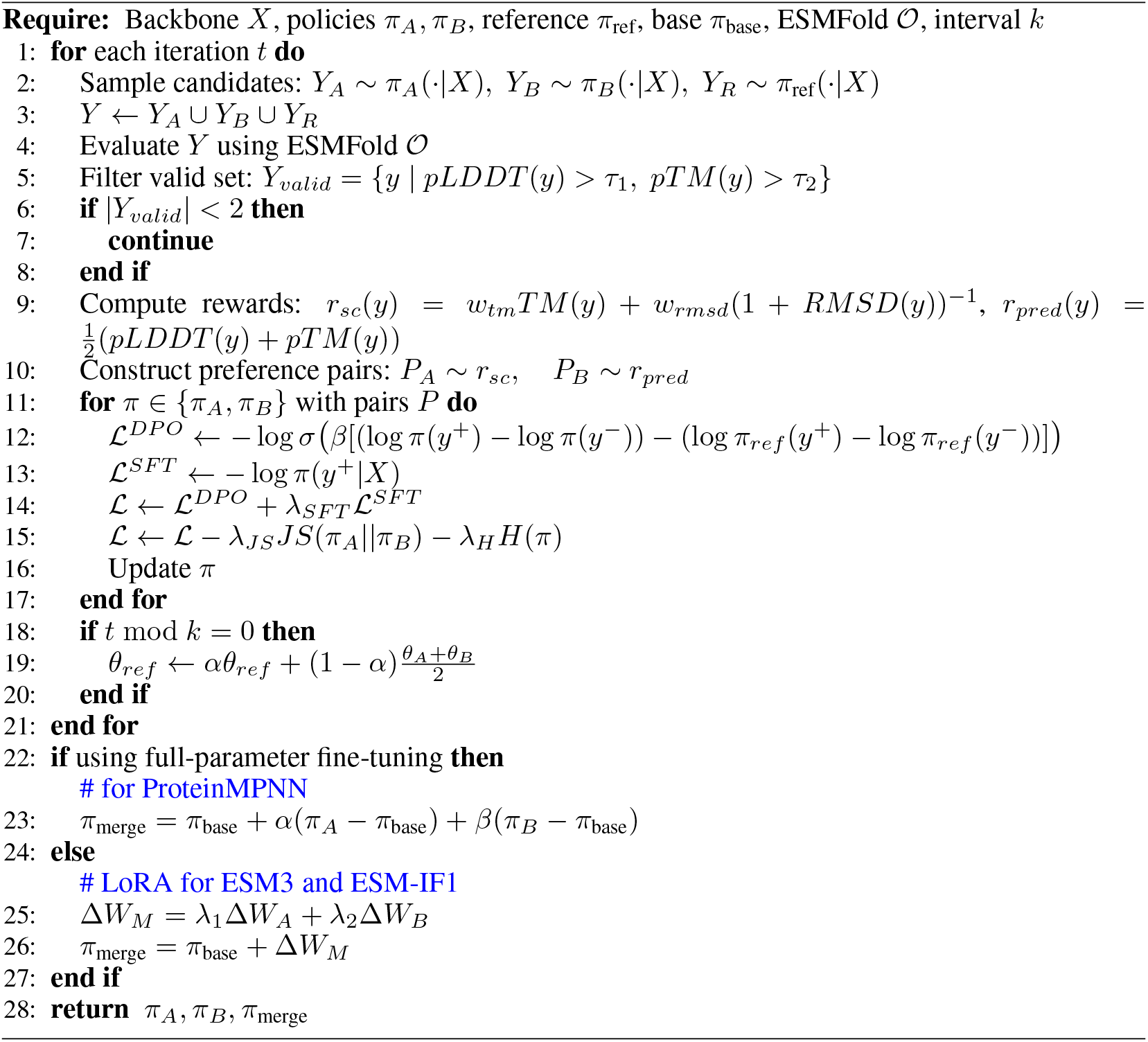

### A.2 Loss Function

The overall training objective consists of four components: a preference-based DPO loss, a supervised fine-tuning (SFT) loss, a diversity-promoting Jensen–Shannon (JS) regularization term, and an entropy bonus.

#### DPO Loss

We adopt Direct Preference Optimization (DPO) to align each policy with high-quality sequences. Given a preference pair (*y*^+^, *y*^−^), the DPO objective encourages the model to assign higher likelihood to the preferred sequence:

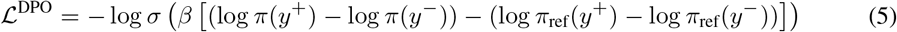

#### SFT Loss

To stabilize training and prevent policy drift, we additionally introduce a supervised fine-tuning (SFT) objective on the preferred samples:

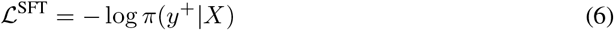

This term ensures that high-quality sequences are directly reinforced, complementing the relative preference signal in DPO.

#### Diversity Regularization

To prevent the two policies from collapsing to similar solutions, we introduce a Jensen–Shannon (JS) divergence penalty:

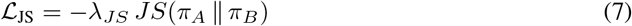

Maximizing the JS divergence encourages the two policies to explore distinct regions of the sequence space, improving coverage of the Pareto frontier.

#### Entropy Bonus

We further include an entropy regularization term to encourage exploration:

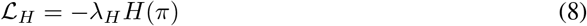

which discourages overly confident predictions and promotes diversity during sampling.

#### Overall Objective

The final loss for each policy is:

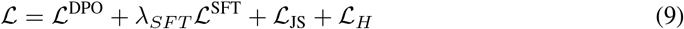

Together, these components balance exploitation of high-quality sequences and exploration of diverse structural solutions, enabling effective multi-objective optimization.

### A.3 Data set

Two widely used datasets in the field of protein IF were employed, namely CATH4.2 and CATH4.3. In addition, this study constructed CAMEO43, Boltzgen-419, and PXDesign-PPI226 from scratch. The detailed components of these datasets are shown in Table 6. Here, we describe the construction of the three newly created test sets in this study.

**Table 6:** The detailed components of datasets in this study.

| Type | Name | # Proteins | Min Len | Max Len | Mean Len | *De novo* Binder |
| --- | --- | --- | --- | --- | --- | --- |
| Training | CATH4.2 | 18,024 | 40 | 192 | 163 | × |
|  | CATH4.3 | 16,699 | 40 | 500 | 228 | × |
| Test | CATH4.2 | 1,120 | 40 | 497 | 161 | × |
|  | CATH4.3 | 1,882 | 40 | 489 | 163 | × |
|  | CAMEO43 | 43 | 33 | 600 | 240 | × |
|  | BoltzGen-419 | 419 | 50 | 500 | 209 | ✓ |
|  | PXDesign-PPI226 | 226 | 51 | 498 | 174 | ✓ |

- **CAMEO43:** We first collected 141 CAMEO targets (January 2025 - April 2025) from THU-ATOM^4^; then, the USalign tool was used to calculate the maximum TM-score of these samples against the CATH training set; finally, only 43 samples with a maximum TM score less than 0.5 were retained.
- **BoltzGen-419:** The original PDB test samples came from EiRA [28] and contained 276 DBPs, 253 RBPs, and 276 PBPs. We removed samples with target lengths > 50 (or < 5) along with those contained non-standard residues. The maximum number of target chains in a single entry is 3. For the remaining targets, BoltzGen was used to generate the *de novo* binder (189 DNA-binding backbones, 77 RNA-binding backbones, and 153 peptide backbones). It is worth noting that no hotspot was specified among these targets. Binder length ratio: [50-200] : [201-400] : [401-500] = 6:3:1.
- **PXDeisgn-PPI226:** To verify the universality of the proposed SSP on *de novo* binders from different generation methods, we used PXDesign to generate protein-binding backbones. The original 398 PDB entries came from the test set of the BoltzGen. We established the following protocol to specify the target chain: **1**. The target chain must have a minimum *Ca* atom distance of less than 8 Å from at least one protein chain; **2**. Residues in contact with other protein chains are designated as hotspots, with a maximum of 10 hotspots (if greater than 10, densely indexed sub-intervals should be selected); **3**. The target chain length should be between 50 and 200; if longer, it should be truncated centered on the hotspots; **4**. The target chain cannot contain any non-standard residues; **5**. A PDB entry produces only one target chain. The interval distribution of the binder length is consistent with that of BoltzGen-419. As a result, 226 protein-binding backbones were produced.

### A.4 Evaluation metrics

Here, we describe the meanings of the metrics used in this study. The structure prediction metrics (pTM, pLDDT, GPDE, and ipTM) come from Protenix [29]; the structure alignment metrics come from USalign (scTM and scRMSD_*Ca*_) and the ESM3 toolkit (scRMSD_*bb*_ and scRMSD_*full*_). All MSAs used for structure prediction were retrieved from UniRef50 and BFD Small databases using `MMseqs2-GPU` [30].

- **pTM:** The global confidence level of the protein structure prediction model for the predicted structure;
- **pLDDT:** Local confidence level of each residue in structure prediction. Protein pLDDT are obtained by averaging the residue pLDDT;
- **GPDE:** Generative pairing distance error. This metric measures the error of the predicted distance for all residue pairs;
- **scTM:** self-consistency TM score (backbone *VS* the predicted structure);
- **scRMSD**_*ca*_ **(scRMSD**_*bb*_, **scRMSD**_*full*_): *Ca* (backbone, full atoms) RMSD score between backbone and predicted structure. Generation with scRMSD_*full*_ < 2Å (scRMSD_*bb*_ for *de novo* binder) is considered a successful design.
- **ipTM:** The predicted interface structure confidence score evaluates interface quality only in the complex structure prediction.

### A.5 Baseline methods

Here, we list the sources of prediction results for all baseline methods. All sequences were sampled at a temperature of 1. Each backbone produced only one sequence for each baseline method.

- **ProteinMPNN:** https://github.com/dauparas/ProteinMPNN. The default model with 0.2 Å noise was employed;
- **ESM-IF1:** https://github.com/facebookresearch/esm/tree/main/examples/inverse_folding;
- **InstructPLM:** https://github.com/Eikor/InstructPLM;
- **InstructPLM-DPO:** https://github.com/Eikor/iplm-rl;
- **MapDiff:** https://github.com/peizhenbai/MapDiff;
- **ADFLIP:** https://github.com/ykiiiiii/ADFLIP;
- **ProteinDPO:** https://github.com/evo-design/protein-dpo;
- **LigandMPNN:** https://github.com/dauparas/LigandMPNN;

### A.6 Molecular dynamics simulation

All MD simulations were performed using `GROMACS` with the `amber14sb_parmbsc1` force field and TIP3P water model. Complexes were solvated in a dodecahedral box with a 1.0 nm buffer and neutralized with 0.15 M NaCl. Energy minimization was conducted using the steepest descent algorithm. The system was then equilibrated under NVT and NPT ensembles (300 K, 1 bar) with positional restraints. Production simulations were carried out for 100 ns with a 2 fs timestep. Long-range electrostatics were treated using PME with a 1.0 nm cutoff, and hydrogen-containing bonds were constrained using LINCS. Temperature and pressure were controlled using the V-rescale thermostat and Parrinello–Rahman barostat, respectively. Trajectories were corrected for periodic boundary conditions and analyzed using RMSD metrics. Binding free energies were estimated using `gmx_MMPBSA`.

### A.7 Implementation detail

We listed the detailed hyperparameters and corresponding descriptions in Table 7. Standard distributed data parallel (DDP) of pytorch was used to implement the SSP algorithm. In the training case of SSP-ESM3 on CATH4.2, we performed 750 policy optimization steps under DDP training. This corresponds to 35,976 backpropagations aggregated across GPUs and 38,408 forward passes. The reference model was updated via EMA every 10 policy optimization steps, resulting in 75 updates. The entire training process lasted approximately 89 hours on 8 RTX Pro 6000 GPUs (96G).

**Table 7:** Hyperparameters list.

| Category | Parameter | Value | Description |
| --- | --- | --- | --- |
| Train | train_epochs | 2 | – |
|  | batch_size | 1 | – |
| | lr | $1 \times 10^{-4}$ | – |
|  | update_every | 12 | Gradient accumulation. The model parameters are updated once every 12 back-propagations. |
|  | ref_update_every | 10 | Reference model updated every 10 policy updates. |
|  | clip_grad_norm | 1.0 | – |
| Base model | ESM3 fine_tune_layer_num | 16 | Last 16 transformer blocks. |
|  | ESM-IF1 fine_tune_layer_num | 4 | Last 4 encoders and decoders. |
|  | ProteinMPNN fine_tune_layer_num | All | Full fine-tuning. |
|  | max_length | 384 | Longer sequences are cropped due to memory limits of ESMFold. |
|  | lora_rank | 8 | - |
|  | lora_alpha | 32 | - |
|  | lora_dropout | 0.1 | - |
| SSP | $\lambda_{js}$ | 0.02 | JS loss weight. |
|  | entropy_bonus_pred | 0.01 | Entropy reward coefficient of *pred*-biased model. |
|  | entropy_bonus_sc | 0.005 | Entropy reward coefficient of *sc*-biased model. |
|  | k_samples | 5 | The number of samples for each policy model. |
|  | k_ref_samples | 5 | The number of samples for reference model. |
|  | temp_policy | 1.0 | Policy model sampling temperature. |
|  | temp_ref | 0.8 | Reference model sampling temperature. |
|  | pair_margin | 0.05 | The minimum allowed reward difference. |
|  | max_pairs | 16 | The maximum allowed number of preference pairs. |
|  | min_plddt | 0.45 | The minimum allowed pLDDT value of samples. |
|  | min_ptm | 0.35 | The minimum allowed pTM value of samples. |
| | dpo_beta | 0.1 | $\beta$ in DPOLoss. |
| | $\lambda_{sft}$ | 0.5 | SFT loss weight. |
|  | ema_decay | 0.995 | EMA update control for the reference model. |
|  | reward_w_tm | 0.4 | The weight of scTM in self-consistent rewards. |
|  | reward_w_rmsd_term | 0.6 | The weight of scRMSD in self-consistent rewards. |
|  | reward_w_plddt | 0.5 | The weight of pLDDT in structure prediction rewards. |
|  | reward_w_ptm | 0.5 | The weight of pTM in structure prediction rewards. |
|  | top_k | 10 | Top-K sampling |
|  | top_p | 0.9 | Top-P sampling |

## Footnotes

3 https://alphafoldserver.com/

4 https://huggingface.co/datasets/THU-ATOM/cameo_data

## Notes

https://github.com/wwzll123/SSP

### Competing Interest Statement

The authors have declared no competing interest.

